# Behavioural foundation of a massive mitochondrial introgression in the fire salamander, *Salamandra salamandra*

**DOI:** 10.1101/2022.08.03.502637

**Authors:** Andrea Chiocchio, Erica de Rysky, Claudio Carere, Giuseppe Nascetti, Roberta Bisconti, Daniele Canestrelli

## Abstract

Patterns of mito-nuclear discordance across secondary contact zones have been reported in a wide range of animal and plant organisms. They consist of a spatial mismatch between nuclear and mitochondrial genomes, in terms of the geographic location and/or extension of the secondary contact zone between distinct evolutionary lineages. Several theoretical and empirical studies have identified massive mitochondrial introgression as the putative source of these mismatches. Yet, we still lack a clear understanding of the potential phenotypic underpinnings of these instances of massive introgression. In this study, we addressed the hypothesis that mtDNA variation across a contact zone could be associated with variation at phenotypic traits affecting dispersal propensity. We analyzed patterns of behavioural and genetic variation across a mtDNA secondary contact zone of the fire salamander *Salamandra salamandra* in central Italy, which is over 600 km displaced from its nuclear counterpart. We found distinct behavioral profiles associated with the two mitotypes co-occurring in the mtDNA secondary contact zone. Counterintuitively, we found a ‘slow-thorough’ dispersal profile associated with the massively introgressed mitotype. This dispersal profile was characterized by shy, less active and less exploratory personality traits, and this pattern was consistent across life-stages and contexts (*i*.*e*., aquatic larvae and terrestrial juveniles). Our results provide experimental evidence supporting the intriguing hypothesis that personality traits associated with distinct mitotypes could promote differential mitochondrial introgression within alternative nuclear backgrounds.

## Introduction

Secondary contact zones are geographic regions where spatially isolated evolutionary lineages meet after range expansion, and they have long been used as natural systems to investigate eco-evolutionary dynamics between and within species (Hewitt 1988). Indeed, when hybridization between divergent lineages occurs, a broad and dynamic range of spatial patterns of genetic variation can arise, leading to a diversity of hybrid zone structures (Barton and Hewitt 1985; Abbott *et al*. 2013). Within this diversity of possible outcomes, considerable attention has been paid to instances of mito-nuclear discordance across contact zones (see Toews and Brelsford 2012 for a review), which consists in geographic location and/or spatial extension of a contact zone between mitochondrial genomes not matching with that defined by nuclear genomes (Toews and Brelsford 2012).

Multiple mechanisms have been proposed to explain patterns of mito-nuclear discordance. Some authors emphasized the role of random processes such as genetic drift and demographic disparities on differential hybrid zone movement (Funk & Omland 2003; Currat *et al*. 2008; Krosby and Rohwer 2009). Other authors emphasized deterministic processes instead, such as sex-biased mating/dispersal or differential fecundity among hybridizing lineages (Petit and Excoffier 2009; Ng and Glor 2011; Rheindt and Edwards 2011). Among the latter, some stressed the role of positive selection and adaptive introgression of mitochondrial genotypes (Hedrick 2013; Bonnet *et al*. 2017; Hill 2019a). Accordingly, massive mitochondrial introgression events would occur if the receiving lineage acquires fitness advantage from the introgressing mitotype, rather than fitness costs by mito-nuclear incompatibility (Dowling *et al*. 2008; Sloan *et al*. 2017; Hill 2019a, b). As a matter of fact, mitochondria are the powerhouse and the master switch of the organism (McKenzie *et al*. 2019), affecting a wide range of phenotypic and life-history traits like respiration, metabolic activities, athletic performances, sexually selected traits, lifespan, and even behaviour (Ballard and Melvin 2010; Breton *et al*. 2014; Hood *et al*. 2018), which hampers the identification of any specific selective advantage associated to cases of mitochondrial introgression.

Dispersal is a key process in ecology and evolutionary biology, as it contributes to shaping spatial patterns of biodiversity and their variation over time (Kokko and López-Sepulcre 2006; Clobert *et al*. 2012; Little *et al*. 2019). Given the key role of mitochondria in energy production and allocation, a causal-effect link between mtDNA variation and energetically demanding behaviours, such those promoting dispersal, in explaining mito-nuclear discordance across secondary contact zones seems especially plausible (Careau *et al*. 2011; Travis *et al*. 2012; Le Galliard *et al*. 2013). Interestingly, experimental evidence reported that mitochondrial introgression can affect animal behaviour, as novel mito-nuclear combinations obtained by artificial hybridization generated more active behavioural profiles (Løvlie *et al*. 2014). Also, personality traits have been shown to contribute to modulate individual dispersal (Bowler and Benton 2005; Cote *et al*. 2010). For example, often fast dispersal profiles are characterized by bold, aggressive, risk-taking, and explorative attitudes (Réale *et al*. 2010; Cote *et al*. 2017). Thus, introgressed mitotypes might carry phenotypic features adaptively linked to increased dispersal capacity, thus providing phenotypic foundations explaining cases of massive mtDNA introgression (Canestrelli *et al*. 2016a, b). However, instances of mito-nuclear discordance in natural populations have been so far mainly investigated by analysing the spatial patterns of genetic variation, while we still lack data for a clear understanding of their phenotypic foundation.

Here, we investigate the phenotypic underpinnings of a striking case of discordance in patterns of geographic variation between mitochondrial and nuclear genes in the fire salamander *Salamandra salamandra*. Recent population genetic studies (Bisconti *et al*. 2018) confirmed the occurrence of two distinct evolutionary lineages along the Italian peninsula, *S. s. salamandra* (the northern lineage) and *S. s. giglioli* (the peninsular lineage; Sillero *et al*. 2014), and outlined the occurrence of a secondary contact between these lineages. However, while nuclear genetic markers located the contact zone in north-western Italy, mtDNA markers spotted it about 600 km to the south, in south-central Italy (Bisconti *et al*. 2018). This geographic pattern was explained by a massive southward introgression of the northern mtDNA lineage into the nuclear background of *S. s. giglioli*, triggered by the post-glacial secondary contact in north-western Italy. Despite mtDNA introgression has been previously reported in salamanders, the geographic extent of this mitonuclear discordance is way more conspicuous (*i*.*e*., Denton *et al*. 2014; Canestrelli *et al*. 2014; Johnson *et al*. 2015). Also, the available data suggest that the mtDNA contact zone in central Italy occurs within a fully “southern” nuclear background (Bisconti *et al*. 2018). Thus, the Italian populations of the fire salamander provides an excellent study system to investigate the possible phenotypic foundation of massive mtDNA introgression. To this aim, we concurrently characterized mitochondrial, nuclear and phenotypic behavioural variation within the fire salamander mtDNA contact zone in south-central Italy. Moreover, to investigate putative carry-over effects among distinct life-stages, phenotypic variation was analysed in the same individuals across the larval and the post-metamorphic life stages, since both these stages - and the associated carry-over effects - can be implicated in the invoked adaptive and non-adaptive processes leading to mito-nuclear discordance.

## Methods

### Study species

The fire salamander *S. salamandra* is an ovoviviparous urodele amphibian (Salamandridae) widespread in temperate and boreal forests of Europe (Sillero *et al*. 2014). Females lay aquatic larvae in small brooks and ponds, and larvae metamorphose after 1-3 months; juveniles have fully terrestrial habit. Sexual maturation occurs after 2-3 years.

Dispersal mainly occurs in larvae and juveniles: larvae actively and passively move across ponds, due to individual resource needs (*i*.*e*., food or refugia, Cecala *et al*. 2009) and drifting, respectively (Thiesmeier and Schuhmacher 1990; Segev and Blaustein 2014), while the metamorphosed juvenile stage represents the highest dispersal phase; adults show strong breeding site fidelity and territorial behaviour (Warburg 1994; Mathis *et al*. 1995; Schulte et al., 2007).

### Sampling and housing

We collected 163 larvae across the secondary contact zone between the two mitochondrial lineages in the Picentini mountains, central Italy (see Figure 1 and Supplementary Table 1). To reduce the probability of kinship among collected individuals, larvae were collected from different breeding sites located few km apart. We monitored the sampling sites since late winter, in order to gain larvae immediately after their deposition. Sampled larvae were immediately transported to the housing facilities in small boxes containing stream water (2 L) and kept in dark and fresh containers. Larvae were then individually housed under controlled conditions (water temperature: 10-12° C; natural photoperiod). We used plastic baskets (10 × 10 cm) as housing containers, with half terracotta saucer as shelter, all set in collective PVC tanks filled with dechlorinated soft water. Larvae were fed *ad libitum* 3 days a week with live *Chironimidae* and *Tubifex*. After metamorphosis, juveniles were individually housed in the same room under controlled conditions (temperature 18-20° C, humidity 65-70%, natural photoperiod). To house juveniles, we used transparent and micro-perforated plastic boxes (11.5 × 11.5 × 6 cm) as terrariums, containing coconut litter, gravel, dechlorinated soft water and a piece of cork as shelter. Juveniles were fed *ad libitum* with small *Acheta domestica* and *Tenebrio molitor*. All the individuals were released to the collection sites after the experiments.

**Figure 1.**
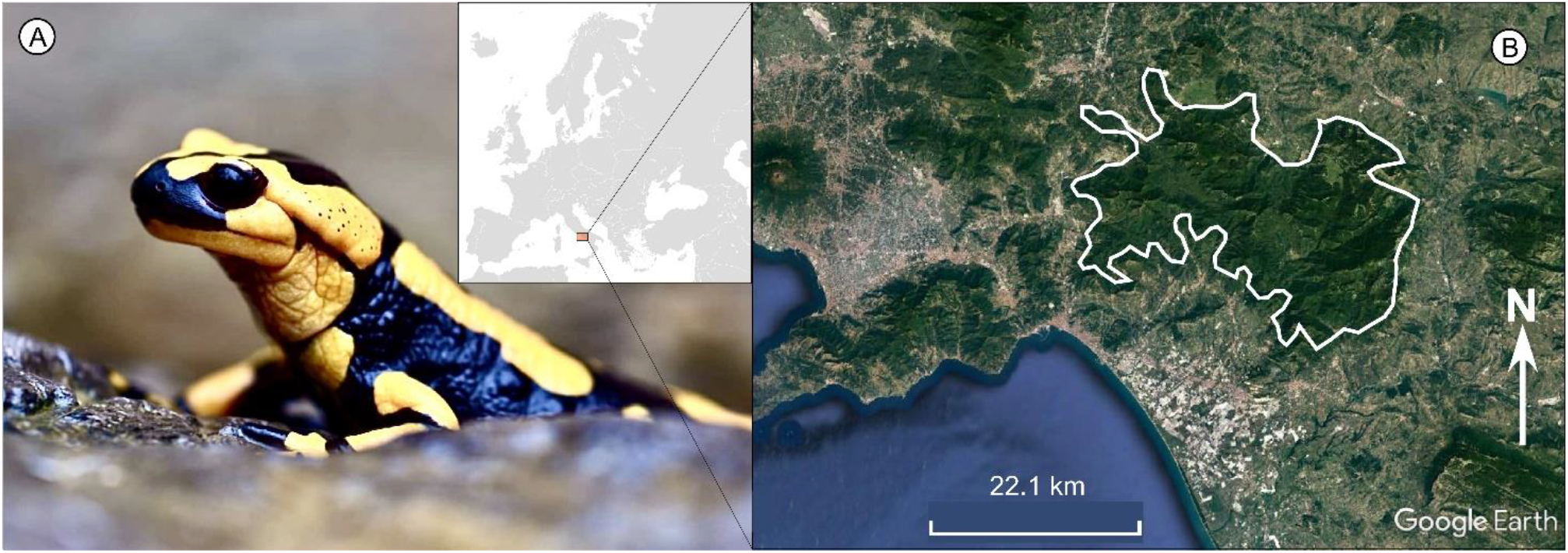
The study species, the fire salamander *Salamandra salamandra* (A), and the study area (B), the Picentini mountain massif (selected area).

All procedures followed the relevant guidelines and regulations for welfare and were approved by the Italian Ministry of Environment (permit number: 0008275.20-04-2018), the Institute for Environmental Protection and Research (#23501, 23-03-2018) and Regione Campania (#0203190, 27-03-2018). Permission to temporarily house amphibians at the University facilities was granted by the Local Health and Veterinary Centre (A.S.L. Tarquinia, license 050VT427).

To obtain individual genotypes, we collected biological tissue for DNA extraction using skin-swabs (Pichlmüller *et al*. 2013). Since the potential influence (if any) of the skin-swabbing procedure on behavioural tests cannot be predicted, this procedure was carried out at the end of the experiments. That is, the entire set of experimental tests was blind to the mtDNA identity of the studied individuals. Before swabbing, individuals were anaesthetized by submersion in a 0.015 m/v solution of Tricaine methanesulfonate (MS222). Then, swabs were stored in ethanol 96 ° at -20°C until DNA extraction.

### Genotyping

DNA extractions were performed using the standard cetyltrimethylammonium bromide protocol (Sambrook *et al*. 1989). mtDNA haplotypes were assessed using the polymerase chain reaction coupled with restriction fragment length polymorphism (PCR-RFLP) diagnostic technique. Diagnostic RFLP profiling was selected by digesting in-silico all the mitochondrial haplotypes from the northern and southern lineages retrieved in Bisconti *et al*. (2018) using REPK (Collins and Rocap 2007), defining two sub-groups (northern and southern haplotypes) and maintaining the default options for defining diagnostic restriction profiles. REPK identified the enzyme *HaeIII* as determining a diagnostic profile in the Cytochrome B (CytB) gene fragment. Then, for all the individuals a 722 bp portion of CytB was amplified by PCR, following the protocol of (34), and subsequently digested by *HaeIII* (Promega) according to the manufacture instructions. Restriction fragments were then separated on 1.5% agarose gel and visualized using UV illumination. Ten specimens from Bisconti *et al*. (2018) representative of the two lineages were used as reference controls.

To assess the effect of local-scale population genetic structure on behavioural variation, and their effect on correlative analyses, all the individuals were genotyped at thirteen microsatellites loci (*SST-A6-I, SST-A6-II, SalE-14, SST-B11, SST-G9, SalE12, SalE2, SST-C3, SalE5, SalE7, Sal29, Sal3, SalE06, SalE8*), which have proved to be informative at fine geographic scale in this species (Lourenco *et al*., 2019). Laboratory procedures for microsatellite amplification followed previously described protocols (Steinfartz et al., 2004; Hendrix et al., 2010). Forward primers were fluorescently labelled, and PCR products were electrophoresed by Macrogen Inc. on an ABI 3730xl genetic analyzer (Applied Biosystems) with a 500-HD size standard. Microsatellite raw data were analyzed using GeneMapper® 4.1. Individuals with more than 50% of missing data were excluded from the subsequent analyses. Micro-Checker 2.2.3 (Van Oosterhout *et al*. 2004) was used to test for the presence of null alleles and large-allele dropout in our data. Allelic frequencies were then computed with GENETIX 4.05 (Belkhir *et al*. 2004) and FSTAT (Goudet 1995) was used to test for deviations from the expected Hardy-Weinberg and linkage equilibria, using the Bonferroni correction for multiple tests.

Populations’ genetic structure within the study area was evaluated using the Bayesian clustering method implemented in STRUCTURE (Pritchard *et al*. 2000). The analysis was carried out using the admixture model, without using sampling location as prior information. Preliminary analyses were conducted to assess model performance, with 80000 steps (the first 20000 discarded as burnin) and 3 replicates for each K value (*i*.*e*., the number of clusters) between 1 and 4. The final analysis contained 10 replicates for each K value, with each run of length 200000 steps, discharging the first 50000 steps as burn-in. The optimal value of K was selected by means of the ln Pr(X|K) method (Pritchard *et al*. 2000) and the Evanno ΔK method (Evanno *et al*. 2005)

### Behavioural assays

The following personality traits were assessed on either larvae and juveniles: spontaneous and exploratory activity, boldness and sheltering behaviour (see Supplementary Table 2 for a summary). Larvae were tested after one week of acclimatization upon arrival; juveniles were tested three months after the metamorphosis. To assess repeatability, each behavioural test was performed twice, after a seven days interval (Bell *et al*. 2009). All individuals were fasted 48 h before the tests. During the tests, room temperature was kept at 20° C and room humidity at 70%. Each behavioural test was recorded using digital cameras (see the next paragraphs for details), and videos were analysed manually with the BORIS software (Friard and Gamba 2016). The tests were set as follow:

#### (i) Spontaneous activity of larvae

The spontaneous activity in the familiar environment (hereafter FE) was measured in the housing container by scan sampling (57). We recorded the activity events with GoPro cameras at predetermined time intervals (2 min/hour, from 10:00 am to 18:00 pm, total 18 min). We also recorded the sheltering behaviour in terms of number of times the animal was inside the shelter with the four paws in.

#### (ii) Boldness and exploratory activity of larvae

The boldness and the exploration were measured in an unfamiliar, novel environment (hereafter NE). Boldness was quantified by measuring the latency to exit from the shelter, and exploration was quantified in terms of time spent in locomotor activity on the total test duration (exploratory activity % on 10 min). NE is assumed to represent a risky situation (Réale *et al*. 2007), thus high activity levels and short latencies were associated with an exploratory and bold personality (Beckmann and Biro 2013). The sheltering behaviour was recorded as in the FE. An individual was considered in immobility if it did not change position for more than 3 seconds. The test arena consisted of a glass tank (24 × 19 cm) filled with dechlorinated soft water (2 L, 12° C), containing a plastic leaf with a stone on the top and a plastic duplicate of salamander larvae, all set at opposite corners between each other. Each larva was gently inserted in a PVC tube as shelter and kept dark for 5 min before the tube opening. Outlier individuals, who have never exited from the shelter or exited immediately (latency < 1 s), were removed from subsequent analyses.

#### (iii) Spontaneous activity of juveniles

The spontaneous activity in the FE was measured in the housing container by scan sampling. We took a picture of the arena each 15 min for 24 consecutive hours (total 96 observations), recording any change of position between each consecutive frame (occurrence of activity events). Using infrared cameras, we also recorded the sheltering behaviour, as defined for larvae. To ensure a clear view of the animals, the coconut litter was substituted with a wet paper towel. Animals were kept in this condition for one week before the test.

#### (iv) Exploratory activity of juvenile

The exploration was measured in the NE. Exploration was quantified in terms of the percentage of time spent in locomotor activity on the total test duration (exploratory activity % on 1 h). An individual was considered not moving at all if it did not changed position for more than 3 seconds. Using infrared cameras, we also recorded the sheltering behaviour, as defined above. The arena set was the same as the FE arena (see above), and – following preliminary observations of juvenile behaviour – the novel situation was considered a sudden change in conditions represented by renewing each paper towel and changing the water container and the shelter before starting the trial.

## Statistical analysis

Individual behavioural consistency was assessed by calculating repeatability estimates on repeated measures for each behavioural descriptor using the mixed models implemented in the “rpt.” function of the “rptR” R package (Nakagawa and Schielzeth, 2010; Stoffel *et al*. 2017). Repeatability coefficients “R”, that is, intraclass correlation coefficients, were calculated as the ratio of between-individual variance to total variance with linear mixed effects models (LMM) for Gaussian distributed data, using individual identity as a random factor and the link-scale approximation of R; activity % in the NE for both larvae and juveniles were square root transformed; the normal distribution of residuals was checked by a Kolmogorov - Smirnov test (see Schielzeth *et al*. 2020). Repeatability coefficients for the latency to exit of larvae in NE and for the sheltering behaviour were estimated by fitting generalized mixed effects models for Poisson distributed data, accounting for over-dispersion (Stoffel *et al*. 2017). The 95% confidence intervals (CI) around repeatability estimates were generated by performing 1,000 parametric bootstrap iterations. Variables were considered ‘highly’ repeatable if R > 0.5 or ‘marginally’ repeatable if R > 0.2 (Bell *et al*. 2009; Aplin *et al*. 2015; Baker *et al*. 2018; Dzieweczynski and Crovo, 2011).

The effect of mitotype in explaining the inter-individual behavioural differences was tested by running generalized linear models (GLMs) implemented in the “lme4” R package. We considered only repeatable behaviours, using the values of the first trial. Each model was run entering the behavioural descriptor as dependent variable and the mitotype as fixed factor. A Poisson distribution with log link function was set for the mobility events of larvae and the sheltering of juveniles in FE; a negative binomial distribution was used for the sheltering of juveniles in NE; a Gaussian distribution with identity link function was set for all the other models.

Finally, we evaluated the correlation between repeatable traits, either within and between life-stages using the Spearman’s rank correlation method implemented in the “spearman.test” function of the “pspearman” R package (Savicky, 2009).

## Results

### Genotyping

According to the PCR-RFLP diagnostic profile produced by the restriction enzyme *HaeIII* on the CytB gene fragments, 81 out 163 individuals showed northern lineage mitotype, and 82 showed the southern lineage mitotype.

The final microsatellite dataset comprised multilocus genotypes for 152 individuals at thirteen loci. Micro-Checker 2.2.3 detected no traces of null alleles and large-allele dropout in our data. There were no significant deviations from Hardy-Weinberg or linkage equilibria after applying the Bonferroni correction for multiple tests. The number of alleles per locus ranged from 2 (*SST-G9*) to 13 (*SalE2*). The analyses conducted in STRUCTURE clearly showed that K=1 best fit the data, revealing no traces of genetic structure in the study area. Indeed, although both the ln Pr(X|K) and Evanno’s methods addressed 3 as the best clustering option, the inspection of the bar plots for all putative K values revealed no traces of biologically meaningful population structures for K>1. Results from the Bayesian clustering analysis performed with STRUCTURE are shown in Supplementary Figure 1.

### Behavioural assays

We characterized the following behavioural descriptors: the spontaneous activity of larvae in the FE, the sheltering behaviour of larvae in the FE, the boldness of larvae in the NE, the exploratory activity of larvae in the NE, the sheltering behaviour of larvae in the NE; the spontaneous activity of juveniles in the FE, the sheltering behaviour of juveniles in the FE, the exploratory activity of juveniles in the NE, the sheltering behaviour of juveniles in the NE.

All but two behavioural descriptors resulted significantly repeatable at the individual level across the two trials (Supplementary Table 3): spontaneous activity of larvae in the FE (R 0.379, p < 0.01), spontaneous activity of juveniles in the FE (R 0.494, p < 0.01), sheltering behaviour of juveniles in FE (R 0.213, p < 0.05), latency to exit from the shelter of larvae in NE (R 0.383, p < 0.01), the exploratory activity of larvae in NE (R 0.340, p < 0.01), the exploratory activity of juveniles in NE (R 0.318, p < 0.01), and the sheltering behaviour of juveniles in NE (R 0.224, p < 0.05). On the contrary, the sheltering behaviour of larvae in both FE and NE resulted not significantly repeatable (p > 0.1).

Results from the GLMs showed substantial behavioural differences among the individuals bearing the two alternative mitotype (Table 1). The effect of mitotype resulted significant for all the tested behavioural traits, except for the exploratory activity of juveniles in the NE, and the sheltering behaviour of juveniles in the NE; the latency to exit from the shelter of larvae in the NE was only marginally significant.

**Table 1.** Outcome of generalized linear models using behavioural traits as dependent variable and the mitotype as fixed factor. FE: familiar environment; NE: novel environment; SE: standard error; significant p-values (p ≤ 0.05) are highlighted in bold.

| Variable | Coefficient | SE | z-value | P-value |
| --- | --- | --- | --- | --- |
| **Spontaneous activity in FE** |  |  |  |  |
| Mobility events of larvae | 0.23605 | 0.04532 | 5.208 | **1.91E-07** |
| Mobility events of juveniles | 7.773 | 2.408 | 3.228 | **1.58E-03** |
| Sheltering of juveniles | -1.5355 | 0.2627 | -5.844 | **5.08E-09** |
| **Exploration and boldness in NE** |  |  |  |  |
| Latency to exit of larvae | -54.31 | 27.65 | -1.964 | **0.05** |
| Activity of larvae | 7.561 | 2.826 | 2.675 | **8.59E-03** |
| Activity of juveniles | 4.867 | 3.365 | 1.446 | 0.15 |
| Sheltering of juveniles | -0.307 | 0.1998 | -1.536 | 0.12 |

Overall, individuals of the northern mitotype took longer time to exit from the shelter and showed lower levels of exploratory activity in the NE than individuals of the southern mitoype. Furthermore, individuals of the northern mitotype showed lower spontaneous activity levels and higher use of the shelter in the FE than individuals of the southern mitotype (see Figure 2 and Figure 3). Interestingly, we found consistent patterns in traits tested for both the life-stages: the northern mitotype showed lower spontaneous activity in the FE and lower exploratory activity in the NE levels, both as larvae and as juveniles.

**Figure 2.**
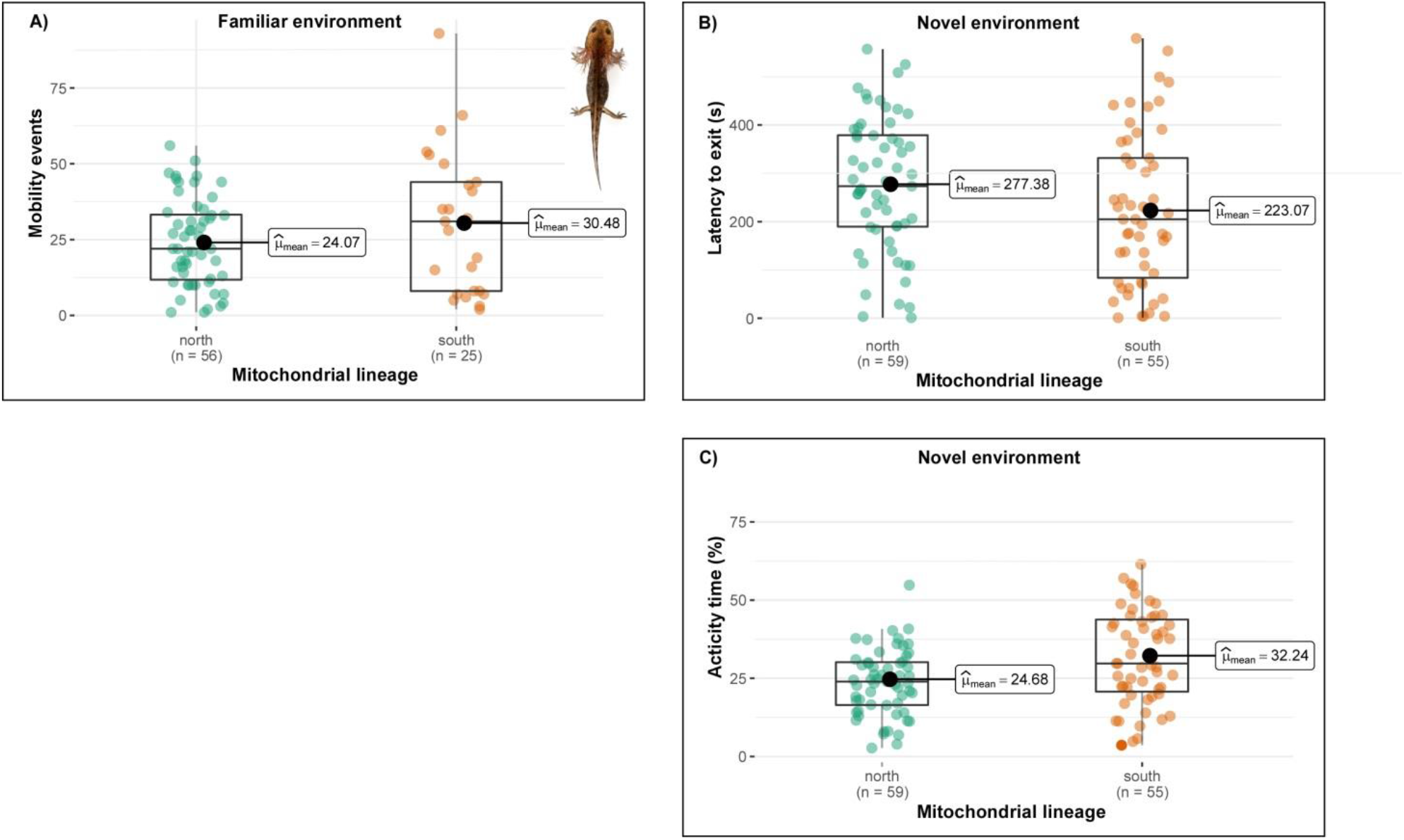
Behavioural traits assessed in fire salamander larvae in familiar and novel environment for the northern and southern mitotypes. (A) Spontaneous activity of larvae in FE expressed as mobility events on 18 min; (B) Latencies to exit from the shelter in NE, in seconds; (C) exploratory activity of larvae in the NE expressed as time spent in activity % on 10 min. Box plots show the mean (inset), the median (central line), the 25 and 75 percentile (box limits), lower and upper bounds (thin lines).

**Figure 3.**
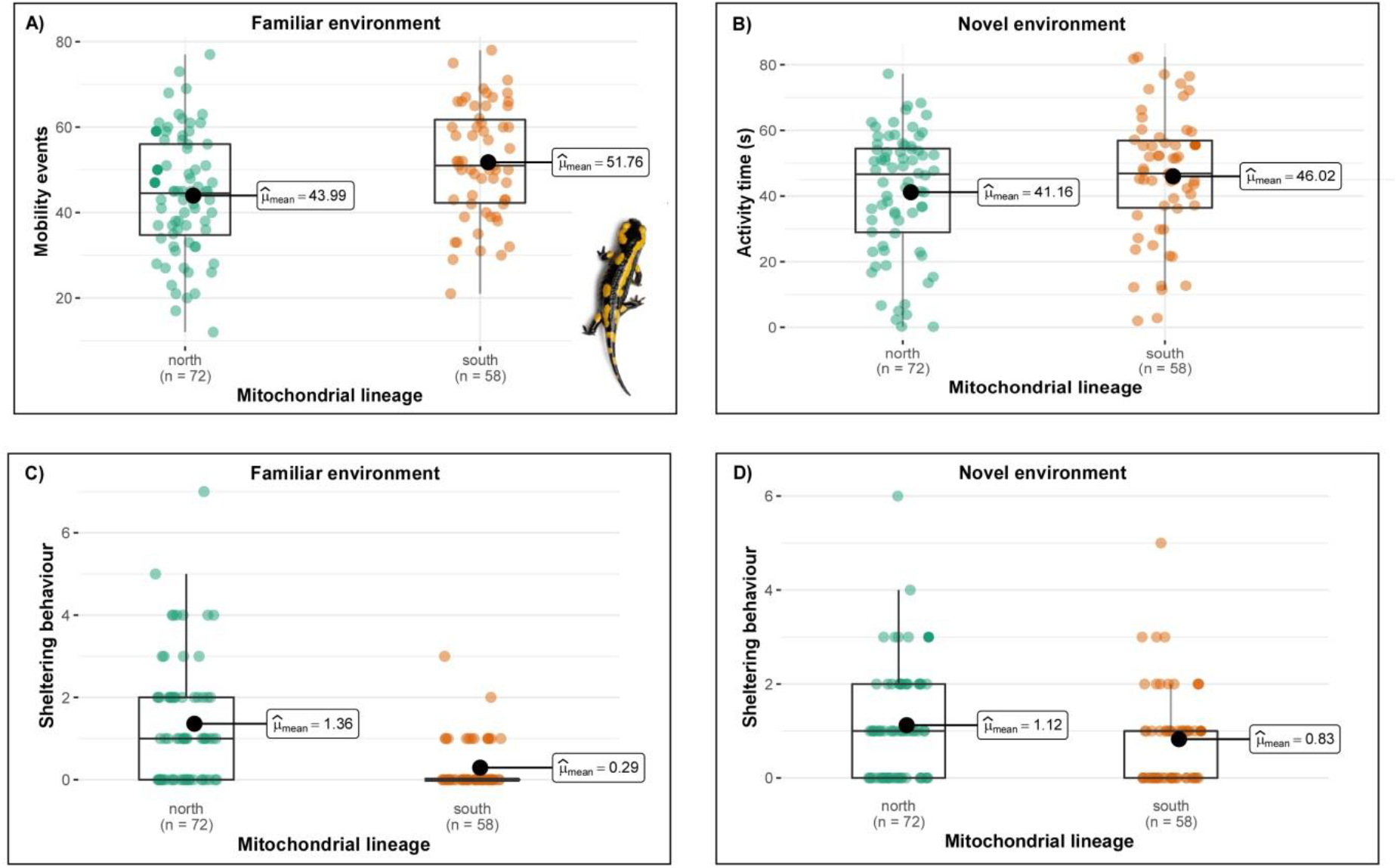
Behavioural traits assessed in fire salamander juveniles in familiar and novel environment for the northern and southern mitotypes. (A) Spontaneous activity of juveniles in FE expressed as mobility events on 24 hours; (B) exploratory activity of juveniles in the NE expressed as % of time spent in activity on 1 h; (C) sheltering behaviour of juveniles in the FE expressed as number of times the individual come back into the shelter; (D) sheltering behaviour of juveniles in the NE expressed as number of times the individual come back into the shelter. Box plots show the mean (inset), the median (central line), the 25 and 75 percentile (box limits), lower and upper bounds (thin lines).

We found no significant correlations among the behavioural traits measured in larvae. On the contrary, we found a negative correlation between the juveniles’ spontaneous activity in the FE and the sheltering behaviour of juveniles in both FE (rS = − 0.236; p = 0.007) and NE (rS − 0.211; p = 0.016). We also found a positive correlation between the spontaneous activity of juveniles in the FE and the exploratory activity of juveniles in the NE (*i*.*e*., less active juveniles used the shelter more than highly active ones, remaining less active in both FE and NE context and *vice versa*; rS 0.211; p = 0.016). Finally, the latency to exit of larvae in the NE was positively correlated with the sheltering behaviour of juveniles in the NE (rS = 0.295; p = 0.006).

## Discussion

Our data support a tight link between mtDNA variation and dispersal behaviour that could explain the observed pattern of mito-nuclear discordance across the secondary contact zone of *S. salamandra* in central Italy. Indeed, we found evidence that two alternative mitotypes of fire salamander spanning the secondary contact zone are associated with two alternative, dispersal-related, behavioural profiles, and that such distinctiveness is consistent across life-stages. In particular, the northern mitotype was associated with a less active and less exploratory behavioural profile, both as larva and juvenile, while the southern mitotype was associated with a more active and exploratory behaviour. Still, such a strong phenotypic differentiation is explained by mitotype in the face of the absence of nuclear genetic differentiation, supporting the hypothesis that the northern mitotype could carry adaptive phenotypic traits, linked to dispersal, driving its rampant introgression (Bisconti *et al*. 2018).

Results from this study showed substantial interindividual variation in dispersal-related behavioural traits of the fire salamander. However, since biologically meaningful variability is conditioned by the consistency of individual patterns, the first important step was to demonstrate statistical repeatability of the traits we focused on (Bell *et al*. 2009). We found individual repeatability and consistency in the behaviour across context. Repeatability and consistency in behavioural responses over time and across contexts have been reported in many taxa, including salamanders, and have been commonly referred to instances of personalities (Gifford *et al*. 2014; Cosentino and Droney 2016; see Kelleher *et al*. 2018 for a review). On the contrary, fewer data are available about consistency across distinct life stages (Koenig and Ousterhout 2018; see Cabrera *et al*. 2021 for a review). We also found consistent patterns in traits tested for both the life-stages, *i*.*e*., the northern mitotype showed lower spontaneous activity and lower exploratory activity both as larvae and as juveniles. Consistency across distinct life stages could represent the outcome of functional links (or constraints) between physiological and behavioural traits, suggesting common underlying mechanisms affecting energy allocation and usage between the larvae and juvenile salamander stages (Biro and Stamps 2010; Gifford *et al*. 2014). This pattern suggests that these personality traits could cluster together with some other life-history trait, such as metabolic activity, body size and life span in “dispersal syndromes”, as observed in many other taxa (Ronce and Clobert 2012; Stevens *et al*. 2014; Cote *et al*. 2017). However, more data from morphological, physiological and performance traits are needed to corroborate this hypothesis.

We found a less active and less exploratory profile associated with the introgressed northern mitotype. This pattern is apparently in contrast with the literature reporting more active and exploratory behavioural profiles at the range edge of expanding populations (Phillips *et al*. 2010; Shine *et al*. 2011; Chuang and Peterson 2016; Gruber *et al*. 2017; Therry *et al*. 2014). However, the higher use of the shelter and the lower exploratory activity in the unfamiliar environment shown by the northern mitotype, in either larvae and juveniles, outline an alternative dispersive profile characterized by long decision-making times in facing the risks of an unfamiliar situation. This in turn outlines a prudent dispersal strategy reflecting a reactive coping style (Koolhaas *et al*. 1999). According to the coping style theory, reactive individuals are “slow-thorough” explorers (*sensu* Koolhaas *et al*. 1999) as they tend to rely more on the information on current environmental conditions, which may take time to acquire, resulting in a cautious, but accurate strategy of exploration (Groothuis and Carere 2005; Réale *et al*. 2010). This strategy could favour the establishment on good feeding ground, due to a more efficient energy balance, and the capacity of performing prolonged efforts (Verbeek *et al*. 1994; Deerenberg *et al*. 1998; Coppens *et al*. 2010), resulting in higher fitness in stressful situations such as the dispersive phase in novel environments (Cockrem 2007; Coppens *et al*. 2010). Thus, the “slow-thorough” dispersal profile associated to the fire salamander northern mitotype might have favoured the northern mitotype introgression because of a higher fitness in recently expanded populations (Clobert *et al*. 2009; Careau *et al*. 2009; Guillette *et al*. 2011). It is worth noting that mitochondria are important modulators of the physiological stress response (Picard *et al*. 2015), whose individual differentiation is a central tenet of the coping style theory. The pattern we found in the two salamander lineages reflecting behavioural profiles similar to the proactive and reactive styles could lead to the hypothesis that mitochondrial function is a major determinant of the different coping styles that could therefore depend upon the type of mitochondrial genome an individual carry. Future research should specifically target mitochondria to understand further the roots of personality differences and stress coping.

### Dispersal syndromes as potential drivers of mitochondrial introgression

Our results support the hypothesis that specific mitotype could carry specific, potentially adaptive, life-history traits linked to the individual dispersal attitude and that such link could lead to mitochondrial genome introgression across a secondary contact zone, irrespective of the nuclear genome background. In the case of the Italian fire salamander, the northern mitotype results linked to specific, potentially adaptive dispersal traits that could have favoured the introgression of its mtDNA into the southern lineage *S. s. giglioli*. The charge to the mitochondrial genotype for the differences in dispersal syndrome’s traits between the two populations of fire salamanders is also supported by lack of any insights for genetic structure at microsatellite nuclear loci in the study area. Thus, the “slow-thorough” behavioural profile associated with the northern mitotypes could be the possible driver of the rampant mitochondrial introgression observed across the secondary contact zone of fire salamander in peninsular Italy. It is worth noting that we assessed the extent of local patterns of genetic structure by employing a limited number of microsatellite loci (see also Bisconti *et al*. (2018), which should not be deemed fully representative of spatial patterns at the level of the entire nuclear genome. Further genetic investigations at the whole genome level should be addressed to investigate the functional genes linked to the molecular pathways driven by the mitochondrial genes, and the geographic structure of their variation across the Italian fire salamander secondary contact zone.

The key role of behavioural polymorphisms in explaining biogeographic patterns is being increasingly recognized, with the hypothesis that divergent evolution of personalities, due to different selection pressure across the range expansion, could lead to a non-random distribution of multiple dispersal strategies - and associated genotypes - among populations (see Canestrelli *et al*. 2016a, b, and references therein). However, from the first idea that individuals characterized by fast-bold behavioural profiles are the main contributors for a successful range expansion, more complex scenarios are emerging. In fact, distinct strategies expressed along the continuums of fast-slow, superficial-thorough, bold-shy, proactive-reactive sets of phenotypic traits within dispersal syndromes can contribute to either the establishment and the reinforcement of new populations in new areas (Bowler and Benton 2005; Duckworth 2008; Clobert *et al*. 2009; Cote *et al*. 2017). Accordingly, the coexistence of those dualisms, and their intermediate modulations, ensure the first spatial exploration and colonization wave in charge to fast-superficial individuals at the expansion front of invading population. In this frame, the northward expansion of the fire salamander southern lineage would be led by fast-superficial (*i*.*e*., bold, Canestrelli *et al*. 2016a) individuals. On the other hand, the subsequent return to the equilibrium in the newly established populations would be in charge of the gene flow of the colonizing populations with the slow-through rears, or by the hybridization of the colonizing populations with the slow-through individuals from the colonized populations, which in turn allow the permeation of slow-through phenotype and genotypes across secondary contact zones (Cobben *et al*. 2015; Canestrelli *et al*. 2016b).

## Acknowledgments

This work was supported by a grant from the Italian Ministry of Education, University and Research (PRIN project 2017KLZ3MA). We thank Michela Paoletti for his precious help during the lab-work, Giacomo Grignani, Giada Spadavecchia and Alessandro Carlini for their support during the field work and the housing activities. We also thank Daniele Delle Monache for his advices on statistics.The authors declare that there is no conflict of interest.

## Author Contributions

D.C. conceived and founded the study. E.d.R., A.C. and R.B. collected the data and performed the experiments. E.d.R. and A.C. performed the data analysis. A.C. and E.d.R. wrote the initial draft of the manuscript, with input from D.C., C.C., R.B. and G.N. All authors edited the final version of the manuscript.

## Data Availability Statement

The datasets generated and/or analysed during the current study are available in the ZENODO repository at the following link https://doi.org/10.5281/zenodo.6799068

## Supporting Information

Additional supporting information may be found online in the Supporting Information section at the end of the article.

Supplementary Figure 1 - Results from the Bayesian clustering analysis performed in STRUCTURE

Supplementary Table 1 - Geographic coordinates of the sampling sites

Supplementary Table 2 - Summary of the tests used to evaluate personality traits

Supplementary Table 3 – Repeatability estimates of the behavioural descriptors

