## Supporting Information for "Behavioural foundation of a massive mitochondrial introgression in the fire salamander, *Salamandra salamandra*"

### Supplementary material

**Supplementary Figure 1** - Results from the Bayesian clustering analysis performed by STRUCTURE on the 13 microsatellite loci (details in the main text; the bar plots show the admixture proportions of each individual for the genetic clusters (K) ranging from 1 to 5.

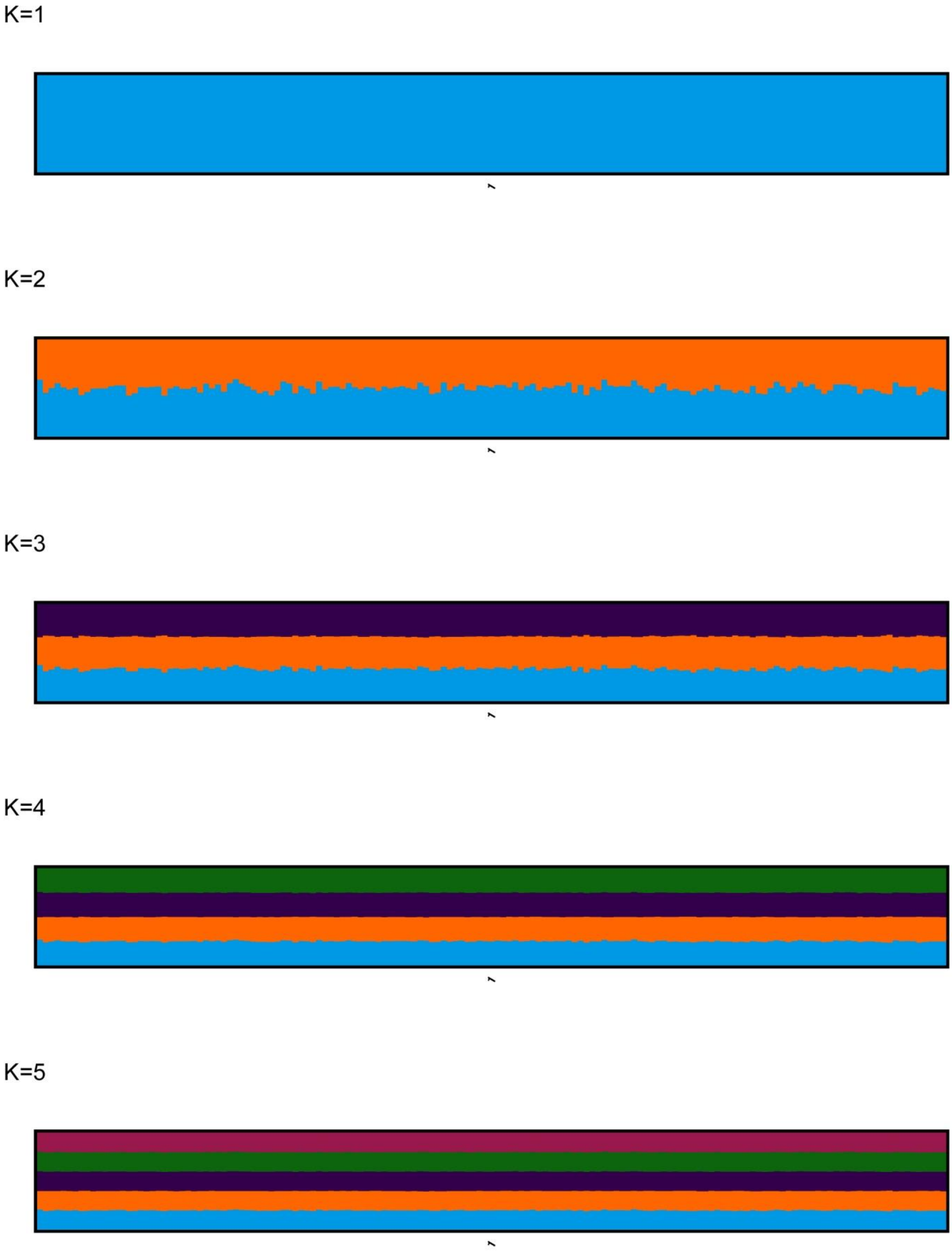

**Supplementary Table 1** - Geographic coordinates of the sampling sites (sexagesimal degrees, DMS).

| Name | Latitude | Longitude |
|---|---|---|
| Montella | 40°45'32.64"N | 15°2'16.38"E |
| Giffoni | 40°46'4.32"N | 14°59'34.14"E |
| | 40°44'33.00" N | 14°59'32.00" E |
| Piani di Verteglia | 40°49'15.72"N | 14°58'32.16"E |
| | 40°48'52.56"N | 14°58'36.48"E |
| Serino | 40°48'5.76"N | 14°53'15.18"E |

**Supplementary Table 2** - Summary of the tests used to evaluate personality traits in larvae and metamorphosed juveniles of the two mitochondrial lineages. Familiar environment (FE), novel environment (NE).

| Test | Life-stage | Duration | Variables | Description |
|---|---|---|---|---|
| **Spontaneous activity in FE** | Larvae | 2 min/h for 9 hrs for a total of 18 min | Mobility events | Number of times the animal makes any locomotor activity (walking and/or swimming and/or change of position) |
| | | | Sheltering | Number of times the animal is inside the shelter with the four paws in |
| | Juveniles | One observation each 15 min, for 24 hrs, for a total of 96 observations | Mobility events | Number of times the animal makes any locomotor activity (walking and/or swimming and/or change of position) |
| | | | Sheltering | Number of times the animal is inside the shelter with the four paws in |
| **Boldness and explorative activity in NE** | Larvae | 10 min | Latency to exit | Time to exit of the shelter with the four paws out |
| | | | Activity % | Percentage of time spent in locomotor activity (walking and/or swimming and/or change of position) |
| | | | Sheltering | Number of times the animal is inside the shelter with the four paws in |
| | Juveniles | 1 h | Activity % | Percentage of time spent in locomotor activity (walking and/or change of position) |
| | | | Sheltering | Number of times the animal is inside the shelter with the four paws in |

**Supplementary Table 3** – Repeatability estimates of the behavioural descriptors of spontaneous activity, boldness and explorative activity for both larvae and juveniles among different trials, obtained by fitting the generalized linear mixed effects models (GLMM) implemented in the "rptR" R package. R: repeatability (i.e. intraclass correlation coefficient); SE, standard error; CI, 95% confidence intervals obtained from 1,000 parametric bootstrap iterations; P: significance, obtained by likelihood ratio tests.

| | **Spontaneous activity in FE** | | | | **Boldness and explorative activity in NE** | | | | |
|---|---|---|---|---|---|---|---|---|---|
| | Larvae | | Juveniles | | Larvae | | | Juveniles | |
| | Mobility events | Sheltering | Mobility events | Sheltering | Latency to exit | Activity % | Sheltering | Activity % | Sheltering |
| R | 0.379 | 0.13 | 0.494 | 0.213 | 0.383 | 0.34 | 0 | 0.318 | 0.224 |
| SE | 0.096 | 0.11 | 0.067 | 0.113 | 0.093 | 0.1 | 0.089 | 0.083 | 0.11 |
| CI | [0.184,0.572] | [0,0.372] | [0.353,0.616] | [0,0.405] | [0.187, 0.548] | [0.124,0.526] | [0,0.297] | [0.157,0.473] | [0,0.395] |
| P [LRT] | 0.000199 | 0.166 | 1.08E-09 | 0.0451 | 0.000168 | 0.00215 | 0.499 | 0.000111 | 0.0312 |
| [Permutation] | 0.002 | 0.189 | 0.001 | 0.069 | 0.12 | 0.002 | 1 | 0.002 | 0.019 |
